# Perturbation variance suppresses error sensitivity in the implicit learning system

**DOI:** 10.1101/2022.09.26.509572

**Authors:** Scott Albert, Reza Shadmehr

## Abstract

When variability is added to a sensorimotor perturbation, total adaptation is impaired. In Albert et al.^1^ we explored this phenomenon, and observed that it is the brain’s implicit, i.e., subconscious learning system that is most affected by perturbation variance. We observed that perturbation variability impaired implicit learning by downregulating its sensitivity to error. Recently, Wang et al.^2^ present an alternate viewpoint: implicit error sensitivity does not change with experience, only the errors observed by the implicit system change. Here we evaluated this alternate view by empirically measuring error sensitivity as a function of error size. We found that perturbation variability strongly downregulates implicit error sensitivity when controlling for error size, consistent with our original results, counter to the inflexible model argued by Wang et al. With that said, a pre-existing relationship between error sensitivity and error magnitude noted by Wang et al. can contribute at least in part to implicit behavior. State-space models that start with this pre-existing error sensitivity curve and then update it with training according to a ‘memory of errors’ most accurately tracked measured behavior.

## Main

Perturbation variability impairs motor learning^3–5^. In Albert et al.^1^ we show that such impairments involve the implicit motor learning system. Our work suggested that suppression in motor learning in response to perturbation variability is consistent with an implicit memory of errors model^6^; namely, with high variance perturbations, error consistency decreases, causing a decline in the implicit system’s sensitivity to error. Wang et al.^2^ propose an opposing viewpoint. In simulation, they argue that suppression in implicit learning could be due not to changes in error sensitivity over time, but to the relationship between error sensitivity and error. They propose a rigid motor correction model, where perturbation variability does not alter the response to error, but simply how these responses are sampled.

The matter at hand, however, is not a model. This debate concerns a fundamental property of the implicit learning system: can the implicit system flexibly alter its response to error? Albert et al. propose that the implicit response to error is modulated by experience. Wang et al. propose that experience plays no role.

These views can be represented at a high-level by the inflexible and flexible models in Fig. 1. The inflexible model (Wang et al.) proposes an identical error sensitivity across the zero and high-variance environments that declines with error (Fig. 1A). Decline in error sensitivity with error causes the motor correction curve (error sensitivity times error size) to exhibit a concave-down shape (Fig. 1B). Perturbation variance alters how this concave-down function is sampled, ultimately suppressing implicit learning (Fig. 1C).

**Figure 1.**
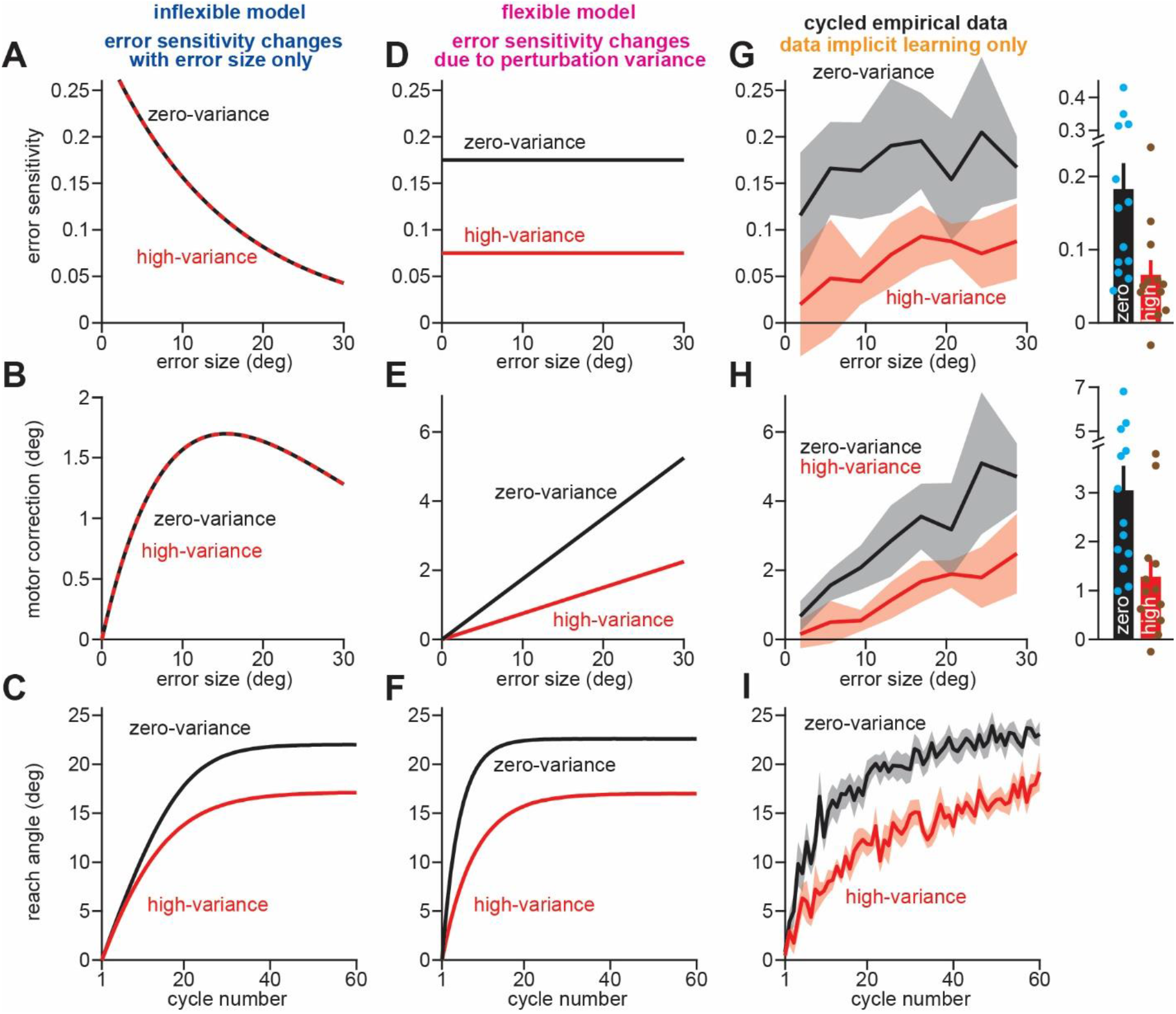
Perturbation variance reduces implicit sensitivity to error. **A**. Error sensitivity in the inflexible (Wang et al.) model: a rigid error sensitivity curve that decreases exponentially with error size but is identical across both the zero and high-variance groups. **B**. The motor correction curve corresponding to the error sensitivity in **A. C**. The average adaptation curve across 100000 simulations using the error sensitivity curve in **A. D**. Error sensitivity in the flexible (Albert et al.) model: high-variance error sensitivity is suppressed by perturbation variance. **E**. The motor correction curve corresponding to the error sensitivity in **D. F**. Same as **C** but for the flexible model in **D. G**. The error sensitivity curve estimated empirically in Exp. 6 in Albert et al.^1^ Here, we cycle-averaged reach angles when calculating error sensitivity to reduce motor noise impacts (see Supplementary Methods). Error sensitivity measures were placed in 3°-wide error bins. At right, error sensitivities were averaged across bins, weighting each bin equally. Dots show individual participants. **H**. Same as in **G** but for the motor correction curves (this is identical to error sensitivity except there is no divisive normalization by error size). **I**. The zero-variance and high-variance learning curves in Exp. 6 in Albert et al. Bars in **G** and **H** show mean value. Lines in **G** and **H** at right show SEM. Lines and shaded error regions in **G, H**, and **I**, show mean ± SEM.

The flexible model (Albert et al.) proposes that error sensitivity is downregulated by perturbation variance (Fig. 1D). Reduction in error sensitivity suppresses the implicit response to error in the high-variance group (Fig. 1E). Finally, this decrease in motor correction lowers overall implicit learning (Fig. 1F).

These simple models highlight a key point: a change in total adaptation across the zero-variance and high-variance groups cannot be used to arbitrate between the opposing viewpoints; both frameworks predict that variance will reduce overall adaptation (compare Figs. 1C and 1F). Instead, we have to measure each condition’s error sensitivity curve and motor correction curve. Here each viewpoint makes a contrasting prediction: the inflexible model (Wang et al.) requires that the response to error is the same across zero-variance and high-variance environments (Figs. 1A,B). The flexible model (Albert et al.) requires that the response to error is altered by perturbation variance (Figs. 1D,E).

In Albert et al. we conducted this analysis (see Fig. 4A and Supplementary Figure 1A in Ref. 4) and observed that the error sensitivity and motor correction curves are suppressed by high perturbation variance. This would invalidate the Wang et al. hypothesis. Our analysis, however, was arguably limited in a couple ways. First, we collapsed across experiments that involved both implicit and explicit learning. This makes it hard to directly see the implicit contributions. Second, we calculated responses to error in wide error bins which limit each curve’s resolution. And third, as noted by Wang et al., motor noise (variability in reaching angle) could alter empirical estimation of learning responses, particularly in the zero-variance condition.

Fortunately, this can all be addressed. First, we can calculate error sensitivity and motor correction solely in Exp. 6 in Albert et al., where reaction time constraints were used to suppress explicit contributions to adaptation. Second, we can calculate the response to error in smaller windows to increase our resolution (we now use 3° error bins as opposed to 8° in Albert et al.). Lastly, we can mitigate how motor noise alters our empirical calculations by using cycle-averaged (i.e., average over all 4 targets in each experiment cycle) as opposed to trial-based measures (the ones simulated in Wang et al.). This approach largely eliminates any effect of motor noise on empirical error response measurements (see Supplementary Figs. 1A-D; also see Section 2 of Supplementary Methods).

Our results are shown in Fig. 1G (error sensitivity) and Fig. 1H (motor correction). Both measures showed an increase in the zero-variance group; perturbation variance reduced the sensitivity to error in the high-variance group (Fig. 1G, t-test on average response, t(23)=2.792, p=0.0104, d=1.118), ultimately reducing their cycle to cycle motor corrections (Fig. 1H, t-test on average response, t(23)=2.798, p=0.0102, d=1.12).

To summarize, we tested how subjects in the zero-variance and high-variance groups responded to error on average across the perturbation period. Our results invalidate the viewpoint proposed by Wang et al. Their hypothesis suggests that the implicit system possesses a rigid and inflexible response to error. This is not true. Variance alters the implicit system’s response to error; consistent with our conclusion in Albert et al., the implicit system flexibly changes its error sensitivity.

We do not mean to suggest that the ideas presented by Wang et al.^2^ make no contribution at all to implicit behavior. Instead, we are more inclined to take a pluralistic view that the implicit system’s error sensitivity is shaped not only by experience (Albert et al.), but also by error magnitude (Wang et al.). These two ideas reinforce one another.

For example, consider the behavior predicted by the Wang et al. error sensitivity model (Fig. 2A). Because their model assumes that the error sensitivity curve is the same across each group and invariant over time, it ultimately underpredicts the initial zero-variance learning rate and overpredicts the initial high-variance learning rate. As a result, this model does not properly capture the timecourse by which the zero-variance and high-variance learning curves diverge (Fig. 2C, compare “data” and “decline only”).

**Figure 2.**
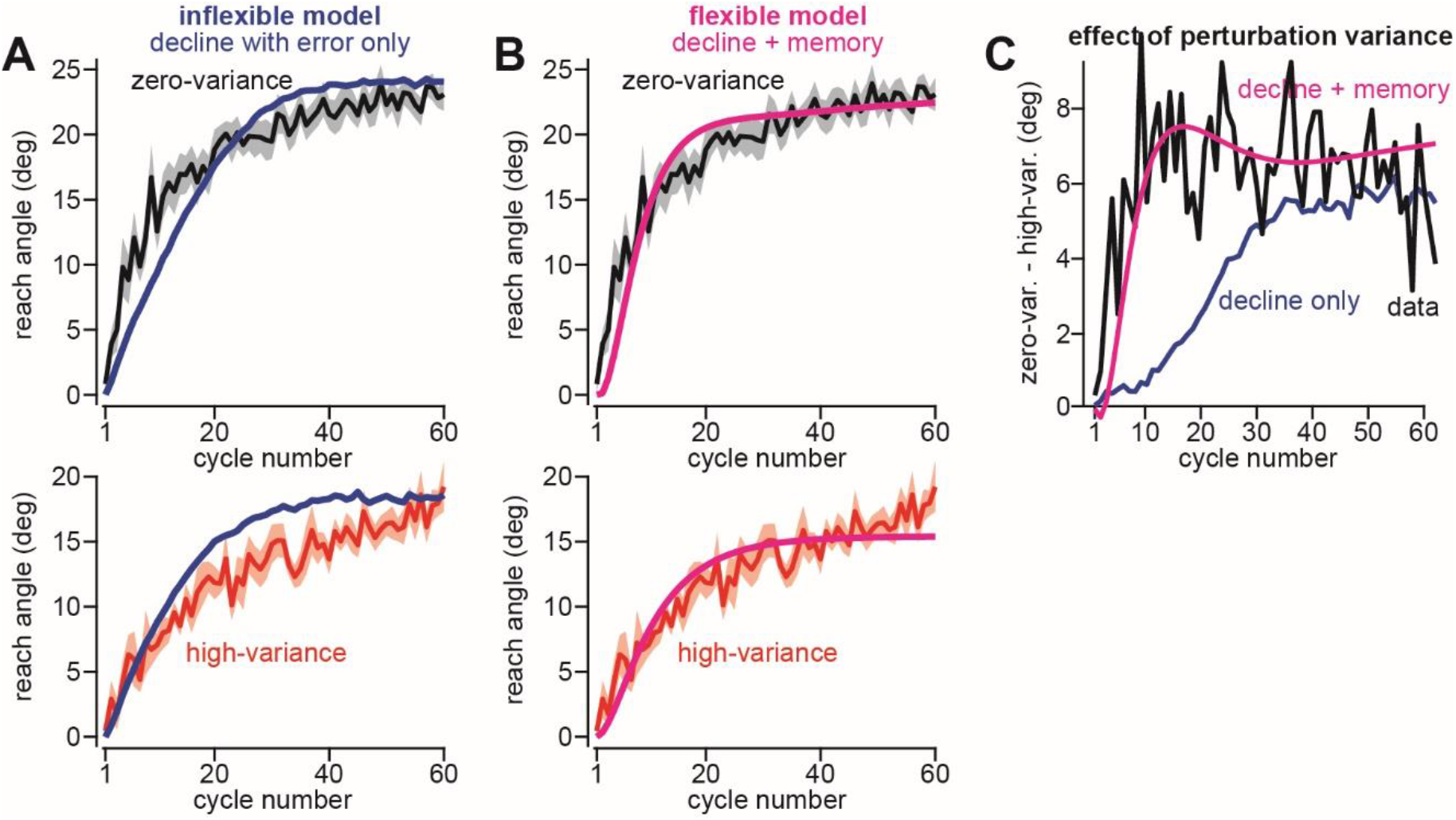
Divergence between zero and high-variance behavior requires error sensitivity modulation. **A**. Comparing zero-variance (top) and high-variance behavior (bottom) to that predicted by the Wang et al. rigid model. Wang et al. predictions are shown in blue and were obtained directly from Fig. 1d in the Wang et al. commentary. The Wang et al. model underpredicts initial learning rate in the zero-variance group (top) but overpredicts the learning rate in the high-variance group (bottom). This indicates that the assumption of a common error sensitivity underestimates zero-variance error sensitivity and overestimates high-variance error sensitivity. **B**. Extending the Wang et al. model with a memory of errors. Here, we simulate a state-space model that combines the Wang et al. and Albert et al. viewpoints. Error sensitivity (see Supplementary Figure 2 and Supplementary Methods) begins with a shared curve that declines with error size, as in the Wang et al. model. However, this curve is modified over time according to the memory of errors model (consistent errors cause a local increase in error sensitivity, inconsistent errors cause a local decrease in error sensitivity). This model produces the behavior shown in magenta. **C**. We subtracted the zero and high-variance learning curves on each cycle. Positive values indicate greater adaptation in the zero-variance group. The ”data” curve shows the timecourse by which the zero-variance and high-variance adaptation curves diverge in Exp. 6 in Albert et al. The blue curve (decline only) shows the inflexible model predictions from Wang et al (obtained from Fig. 1d in Wang et al.). Last, the magenta “decline + memory” curve shows the behavior predicted by the model in **B**. Error bars in **A** and **B** show the mean ± SEM.

These limitations implicate a key implicit learning property: error sensitivity between variance conditions must change over time. To show this, we considered a holistic model in which error sensitivity starts as a curve that declines with error size (Wang et al.) but changes over time due to error consistency according to the memory of errors model (Albert et al.). This model closely tracked the true behavior (Fig. 2B) and predicted the divergence between zero-variance and high-variance learning with high accuracy (Fig. 2C, compare “data” and “decline + memory”).

In summary, the viewpoint proposed by Wang et al.^2^ can be directly tested in the data. The empirical error sensitivity and motor correction curves (Figs. 1G,H) invalidate their premise, and show unequivocally that perturbation variance changes the implicit system’s sensitivity to error. Combining the memory of errors model with the Wang et al. viewpoint provides a closer match to behavior, suggesting that error sensitivity changes with both error (Wang et al.) and time (Albert et al.). Future experiments could test the relative contribution each makes to behavior.

We see in the field a controversy about whether the implicit system has a flexible response to error. Albert et al. provides one piece of evidence that implicit error sensitivity changes, an implicit phenomenon that also appears in other scenarios: altering the cost of our errors^7^, altering uncertainty about error^8^, changing task outcome signals^9,10^, revisiting a past perturbation^11,12^. As the field moves forward, we think it will prove limiting to presume rigidity in implicit learning properties, rather than consider the constellation of factors that might modulate implicit behavior across different contexts. For example, the often-cited rigidity that implicit learning exhibits in its saturation to perturbation size^13,14^, is more likely a symptom of the learning environment it is placed in^13,14^ or the explicit strategies it competes with^15–17^, than an immutable implicit learning phenotype (see Ref. 10). All in all, like the brain, implicit adaptation is complicated, and we have only begun to uncover its code.

---

**Supplementary Figure 1.**
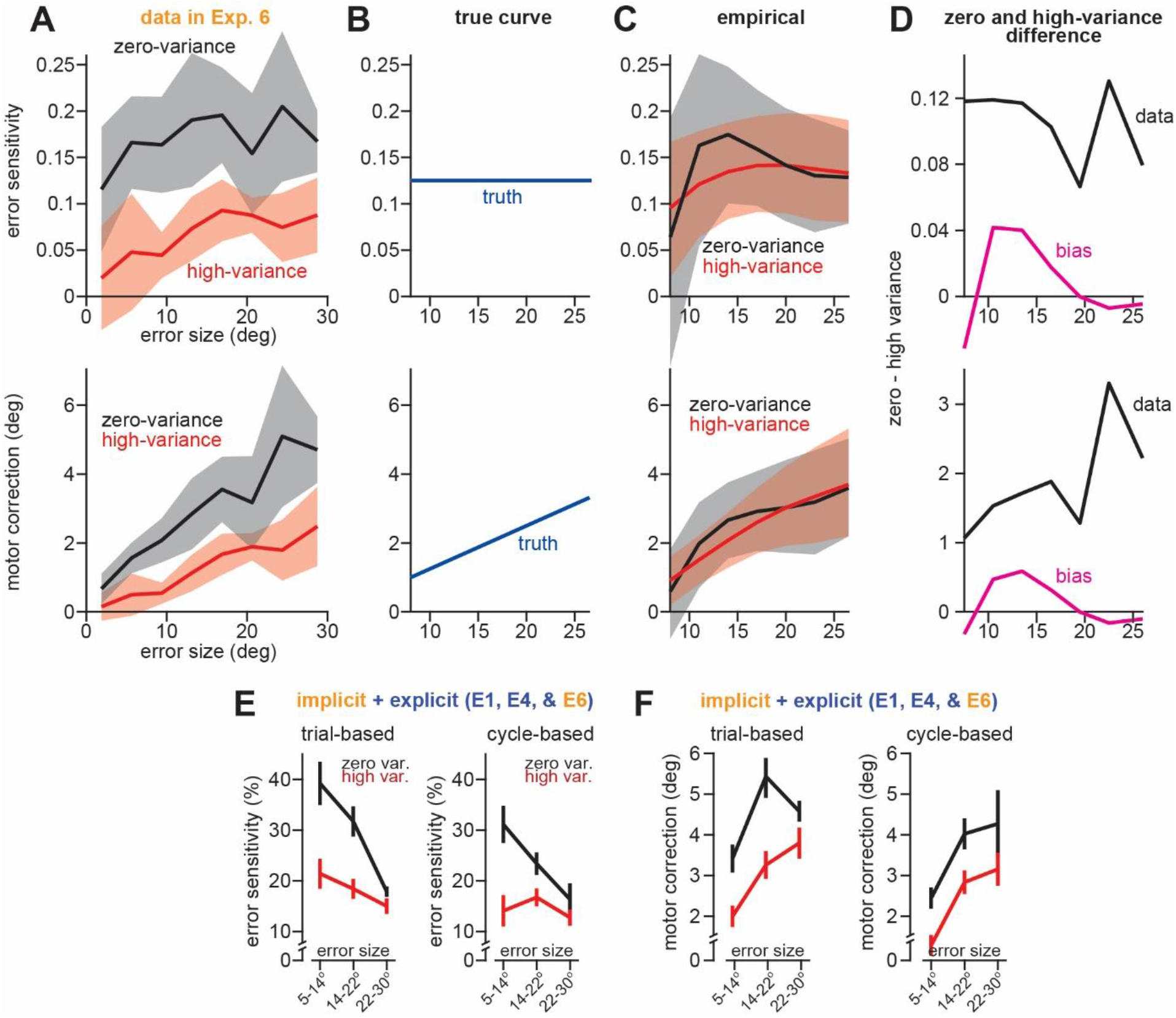
Cycle-averaged reach measures minimize bias. **A**. Here the empirical error sensitivity (top) and motor correction curves (bottom) in Figs. 1G and 1H are reproduced. We want to know the extent to which the motor noise in reaching behavior might contribute to the divergence between the experimental groups. **B**. We used a nominal error sensitivity of 0.125 to simulate participant behavior (top). The corresponding motor correction curve is shown at bottom. Note, in simulation we used trial-by-trials noise levels estimated in the data (σ_y_=4° and σ_x_=0.8°, see Supplementary Methods). To reproduce subject-to-subject variability in asymptotic performance, the simulated error sensitivity was sampled from a uniform distribution whose mean value was 0.125. **C**. Here the error sensitivity and motor correction curves were calculated in simulation, using the same procedure as the actual data in **A**. Solid lines denote the average curve over 100000 simulations. The shaded areas represent 95% confidence on the mean. **D**. The mean bias between the zero-variance and high-variance curves in **C** (the magenta “bias” curve) is compared to the true difference in zero-variance and high-variance error responses in **A** (black “data” curve). The substantial mismatch between the data and bias curves indicates motor noise cannot produce the difference between the error sensitivity and motor correction patterns we estimated empirically in the data. **E**,**F**. In Albert et al., error sensitivity and motor correction curves were calculated with trial-by-trial reach angles (see Fig. 4A and Supplementary Fig. 1A primary figures in Ref. 4). Motor noise can alter error sensitivity measurements using single trial reach angles. We recalculated the error response curves using cycle-averaged reach angles to corroborate our findings. The trial-based error sensitivity and motor correction curves originally reported in Albert et al. are shown at left in **E** and **F**. The same curves are recalculated at right using cycle-averaged reach angles.

**Supplementary Figure 2.**
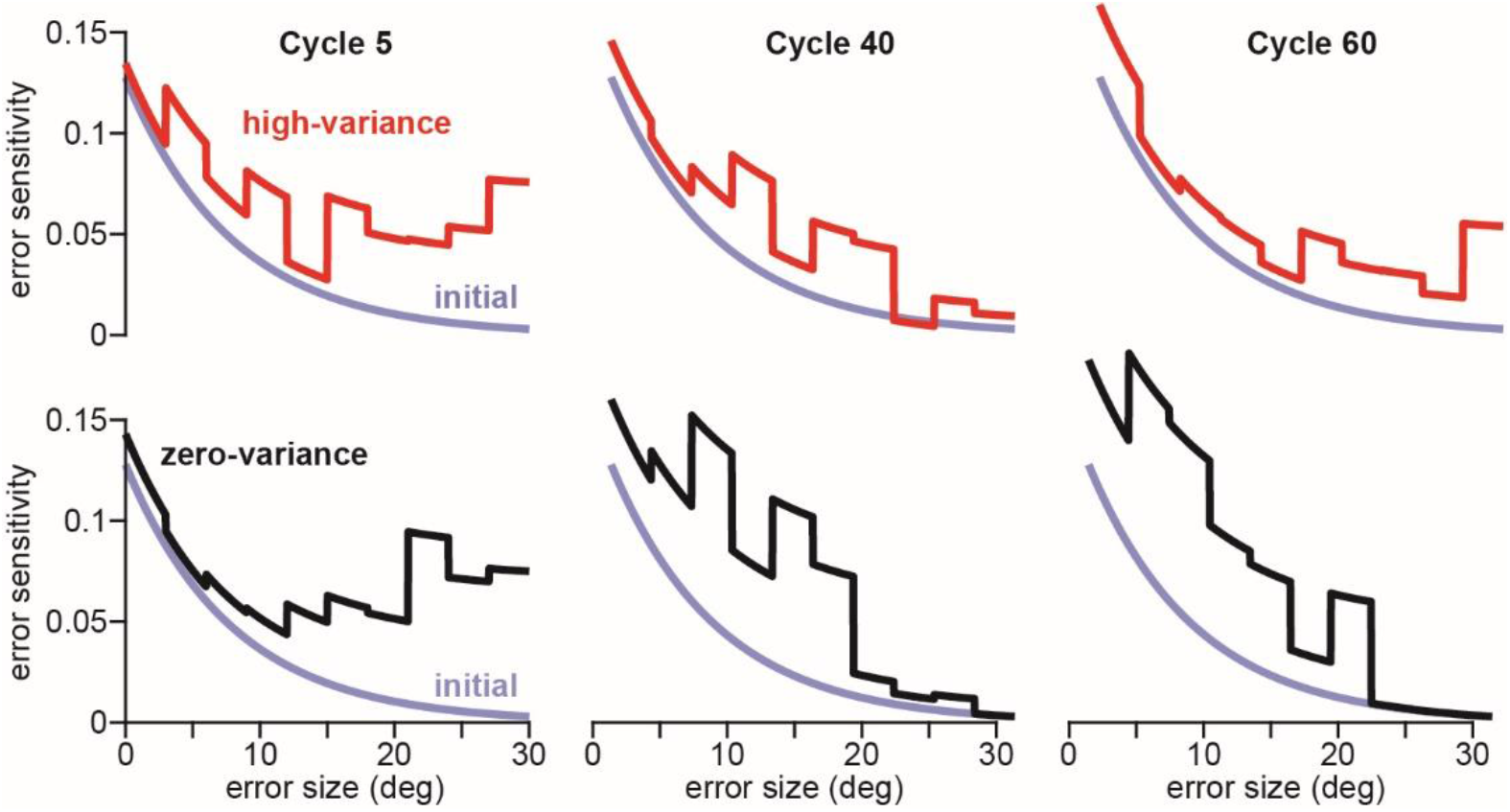
Error sensitivity modulation due to a memory of errors and error size. Here, error sensitivity curves corresponding to the simulations in Fig. 2B are provided (the flexible “decline + memory” model). To reiterate, in this model error sensitivity has a baseline shape by which it decays with error size (this is the property described by Wang et al.). Here, we extended this error sensitivity behavior with a memory of errors. When a consistent pair of errors are experienced (same sign) error sensitivity locally increases. The opposite occurs when the errors are not consistent (opposite sign). Here we show the error sensitivity patterns that emerge early and late during adaptation. Columns 1-3 from left to right correspond to Cycles 5, 40, and 60. These predictions are made using the memory of errors model operating on the errors actually experienced by the 13 participants in the zero-variance group and the 12 participants in the high-variance see (see Supplementary Methods). The top row shows predictions for the high-variance condition. The bottom row shows the zero-variance condition. Changes in the error sensitivity curves are shown in the bolded lines. The faded “initial” lines show the initial error sensitivity curve on Cycle 1; this is identical across the zero-variance and high-variance groups.

## Methods

Here we describe the analyses reported in the main text as well as supplemental calculations that support our conclusions.

### Section 1. Flexible and inflexible implicit learning models

Albert et al.^1^ and the commentary by Wang et al.^2^, possess two opposing viewpoints. Albert et al. suggests that error sensitivity is altered by perturbation variance, which then suppresses high-variance adaptation. Wang et al. presumes variance does not alter error sensitivity. In their view, suppression in high-variance adaptation is due to the way error sensitivity declines with error size.

At its core, the Albert et al. view suggests error sensitivity is truly elevated in the zero-variance group. In this model, implicit learning has a flexible error sensitivity. At its core, the Wang et al. view suggests that error sensitivity is identical in the zero-variance and high-variance groups. It abides by an inflexible, rigid motor correction curve.

These hypotheses are represented at a high-level in Fig. 1. We term the Albert et al. model, the “flexible” model. We term the Wang et al. model, the “inflexible” model. Each model is represented very simply in Fig. 1. To represent the flexible model, the suppression in high-variance learning is entirely attributed to suppression in error sensitivity (and no decline with error size). Thus, in Fig. 1D, we set error sensitivity to 0.075 and 0.175 in the high-variance and zero-variance conditions, respectively. We selected these values so as to achieve approximately the correct total adaptation levels in both groups (see Fig. 1I). The motor correction curves that correspond to these error sensitivities are shown in Fig. 1E. These are calculated as error sensitivity multiplied by error size. In Fig. 1F, we simulate the learning curve that corresponds to the error sensitivities in Fig. 1D. The inflexible model (Figs. 1A-C) uses the same basic process, only the error sensitivity in Fig. 1A is altered. Here the zero-variance and high-variance groups have the same sensitivity to error. However, this error sensitivity declines according to an exponential function: *b* = 0.3 exp(−0.065*e*) where *b* is error sensitivity and *e* is error magnitude (i.e., the absolute value of error).

Lastly, the simulated learning curves in Figs. 1C and 1F are the average response over 100000 simulations. In these simulations, we use a state-space learning model. The structure mirrored Exp. 6 (limited reaction time task) in Albert et al. There were 240 adaptation trials. These trials occurred in cycles where 4 targets were visited once each. This correspond to 60 cycles total.

In the state-space model, each target has its own adaptive state, and learning does not generalize across targets because they are separated by 90°. When target *t* is visited on trial *n*, its state experiences error-based learning according to:

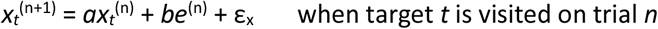

Here, ε_x_ is a normal random variable with mean 0°, and standard deviation σ_x_. For the 3 targets that were not visited on trial *n*, they simply exhibit decay:

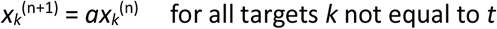

Note that the retention factor is a trial-by-trial retention factor. To obtain this we need to raise the cycle-based retention factor (0.945) to 0.25. This yields *a* = 0.986 on a trial-by-trial basis. This value was selected by calculating the average retention factor during a no-feedback period at the end of Exp. 6 (see Albert et al. for a description on the retention factor estimation procedure). The *b* term represents error sensitivity and was set based on the curves outlined above in the flexible and inflexible model descriptions.

Lastly, the motor command produced on trial *n* is driven by the adaptive state that corresponds to target *t* (the target that is visited on trial *n*). This gives:

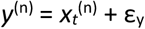

Here, ε_y_ is a normal random variable with mean 0°, and standard deviation σ_y_. This constitutes the model that we used in our simulations.

It is important to select accurate σ_x_ and σ_y_ values, especially the latter as this represents motor noise. As described below, motor noise alters empirical estimation of error sensitivity. To estimate motor noise in the data, we used a process suggested by Wang et al. in pre-submission correspondence. To estimate the σ_y_ we calculated the lag-1 autocorrelation in cycle-averaged reach angles. To ensure outlier reach trials did not unfairly bias the result, we removed trials that deviated by more than 14.5° from an exponential curve fit to the data (14.5° threshold was selected so as to target only the left and right 2.5% tails in the residuals distribution). This outlier removal yielded a final median lag-1 autocorrelation of 0.7364.

The next step was to determine the motor noise level that matched this autocorrelation. To infer this, we simulated the state-space model above and varied σ_y_. For each value, we calculated the corresponding lag-1 autocorrelation. Through this process, we determine an optimal σ_y_ value of 4°. Note that this value is conservative, exceeding other measurements in the literature: e.g., 2.5-3.5° Therrien et al.^18^

Lastly, we also required a value of σ_x_. For this, we also used a suggestion made by Wang et al. during pre-submission correspondence: that state noise is about 20% the size of motor variability^19^. Thus, 20% of our σ_y_ estimate was 0.8°. This is the value used for σ_x_.

### Section 2. Error sensitivity and motor correction curve estimation

The most important analyses in this response are shown in Figs. 1G and 1H. These are the error sensitivity and motor correction curves estimated empirically using the reach angles in Fig. 1I (to reduce estimation noise, outlier trials were removed using a standard 1.5 inter-quartile range benchmark; this is reflected in the reach angles in Fig. 1I).

The error sensitivity curve was calculated using:

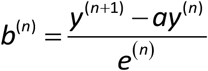

Here *b*^(n)^ is the error sensitivity on cycle *n*. The *y*^(n)^ term is the cycle-averaged reach angle on cycle *n*. The *e*^(n)^ is the cycle-averaged error on cycle *n*. The use of cycle-averaged measures is important here (described further below). The cycle-average simply means averaging over the 4 distinct targets that were presented once each during each experiment cycle. For the motor correction curve, the formula above is used, but change in reach angle is not divisively normalized by error. Because cycle-averaged quantities were used, note the *a* term is the cycle-averaged retention factor.

In Albert et al., we used a similar process to measure error sensitivity, except trial-based measures were used. There error sensitivity and motor correction was calculated within 8° error size windows. To increase resolution, we lowered this bin size to 3° in this response. In other words, the error sensitivities and motor corrections calculated above, were sorted into bins by the corresponding error. The bins were: 5°-8°, 8°-11°, 11°-14°, 14°-17°, 17°-20°, 20°-23°, 23°-26°, 26°-30°. We then averaged the values in each bin across all cycles. Figs. 1G and 1H show the mean and corresponding SEM within each error bin, in the zero and high-variance groups. At right in Figs. 1G and 1H we compared the error sensitivity and motor correction across the zero-variance and high-variance groups. To do this, we simply took the underlying participant curves used to generate Figs. 1G and 1H and calculated the average, weighting each error bin equally. We then compared the zero-variance and high-variance participants using a two-sample t-test.

As alluded to in the main text, and also by Wang et al., empirical error response measures are altered by motor noise. To very briefly explain, during adaptation there is a relationship between reach angles, error size, and motor noise. This relationship is stronger in the zero-variance condition, due to a closer relation between reach angle and error size. This relationship can ultimately bias error sensitivity “upwards” and also “downwards”.

Wang et al., provided some simulations to illustrate this effect. There are two critical things to note here. First, their simulations use trial-based measures, not cycle-averaged measures. This is critical. Averaging reach angles in a cycle reduces overall noise level (it essentially halves it because each cycle has 4 trials). Second, their simulated “subjects” all have the same learning parameters (i.e., *a* and *b*). This is important as well. Across-subject variability in adaptation weakens the population-level relationship between error size and reach angle, ultimately reducing empirical bias. Thus, not including such variation, overestimates the extent to which motor variation will alter error response measures in the actual participant data.

We conducted analyses to ensure the technique (i.e., cycle-averaging) would make our motor correction and error sensitivity calculations robust to motor noise. To do this, we used a method similar to Wang et al. We simulated behavior according to the state-space model in Section 1 above. We set motor noise (σ_y_) equal to 4° and process noise (σ_x_) to 0.8° as described above. For simulated participants we sampled the trial-based retention factor from a uniform distribution between 0.98 and 0.99. Lastly, we used “flat” error sensitivity functions to more easily isolate any bias in error sensitivity. We used a nominal error sensitivity of 0.125. We used this value as it straddled the mean error sensitivity across the zero and high-variance groups (see Fig. 1G). Other choices yield qualitatively similar results. To simulate inter-subject variability, we sampled the error sensitivity from a uniform distribution between 0.045 and 0.205. We used this range so as to match subject-to-subject variance in total adaptation observed in the zero-variance group (with the simulated range we obtained a value of 3.26°, in the data it is 3.29°). Critically, for *a* and *b*, the same parameter distributions were used in the zero and high-variance groups. The goal is to determine whether there may be a bias in estimating error sensitivity between the groups, when it is truly identical.

Finally, we calculated the empirical error sensitivity curves in the simulated data, using the same equations outlined above (the ones used in Figs. 1G and 1H). Ideally, if this empirical estimation is unbiased, the zero and high-variance groups would be identical and match the true curves. Supplementary Fig. 1B shows the true error sensitivity and motor correction curves (i.e., the curves used in the simulation). Supplementary Fig. 1C shows the mean values obtained empirically across 100000 simulations. Error bars show the 95% CI on the mean estimated using bootstrapping; to match the experimental data, 13 (zero-variance) or 12 (high-variance) subjects were sampled from the distribution, and then averaged in each bootstrap. Note, the response in the smallest bin (5°-8°) is omitted, because the nominal 0.125 error sensitivity yields total asymptotic errors above this error range, producing negative error sensitivity values in the zero-variance group (this issue is mitigated in the actual zero-variance data, because zero-variance error sensitivity is higher than 0.125). The mean bias between the zero and high-variance groups (i.e., an offset between the curves in Supplementary Fig. 1C) is quantified in Supplementary Fig. 1D. Here we subtract the mean curves across the zero-variance and high-variance groups and compare that to the difference seen in the actual data (the offset between the black and red lines in Figs. 1G and 1H, reproduced in Supplementary Fig. 1A).

Supplementary Figs. 1C and 1D show that cycle-averaging reach angles, substantially reduces any bias in the empirical error sensitivity and motor correction measures. The average difference across the zero and high-variance curves in Figs. 1G and 1H was an order of magnitude larger than what would occur based on motor noise bias alone. This is despite using very conservative motor noise estimates in simulation, (σ_y_ equal to 4°) that match or exceed the upper bounds observed across young and old adults in the literature (see Ref. 17). In sum, these simulations unequivocally demonstrate that the differences in error sensitivity and motor correction observed in Figs. 1G and 1H reflect a true suppression in the high-variance group’s response to error. As we explained in the main text, this invalidates the Wang et al. view; perturbation variance directly modulates error sensitivity in the implicit system.

Lastly, we re-analyzed data in Albert et al., where error responses were calculated in different error size bins (note the point error sensitivity estimates in Fig. 3 in Ref. 4 (average error sensitivity across all trials), will not exhibit bias; this only emerges when trying to resolve error sensitivity as a function of error size). Note, these data were reported in Fig. 4A and Supplementary Figure 1A in Ref. 4. In our new analysis, we cycle-averaged reach angles prior to measuring error sensitivity and motor correction. The cycle-based (re-analysis) and trial-based (Ref. 4) results are in Supplementary Fig. 1E and 1F in this response. While cycle-averaging reach angles caused a reduction in overall error sensitivity estimates, the same divergence between the zero and high-variance data remained; thus, our conclusions in Albert et al. were also not altered by motor variability.

## 3. Memory of errors model

In the main text, we describe issues with the assumption in the Wang et al. model that the zero-variance and high-variance groups have identical error sensitivities. This causes their model to underpredict initial learning rates in the zero-variance group and overpredict them in the high-variance group. The key insight here is that while error sensitivity may start the same in both variance conditions, it does not stay identical and changes according to the environmental statistics.

Instead of treating the Wang et al. and the Albert et al. models as either-or frameworks, we hypothesized that more accurate predictions would be obtained by combining the error sensitivity properties described by Wang et al. (error sensitivity declines with error size) with the error sensitivity properties argued in the Albert et al. memory of errors (error sensitivity is altered over time in accordance with error consistency).

We created a model that integrated both hypotheses. This model is virtually the same as that reported in our original paper (see Ref. 4), except we treat baseline error sensitivity as a curve that declines with error size (in the original paper, initial error sensitivity was constant across error size, for the sake of simplicity). For convenience, we will reiterate the most important model components here. Our updated memory of errors model uses the state-space model in Section 1 but treats error sensitivity as a function that updates over time (i.e., the *b* is not constant in the equations).

Here, we imagined that error sensitivity has same baseline state *b*_*0*_ which it tends to decay towards. This baseline state was treated as a decaying function of error, consistent with the Wang et al. hypothesis. We set baseline error sensitivity according to:

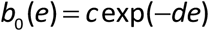

Here *c* and *d* are unknown parameters that describe how baseline error sensitivity *b*_0_ decays with the error magnitude *e*. Error sensitivity on any given trial is determined by past modifications to *b*_0_. The relationship between *b* and *b*_0_ is given by:

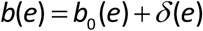

Here the *e* in parentheses is meant to denote that all quantities are a function of error. The δ parameter is the modulation to baseline error sensitivity. It is determined by a memory of errors (with learning and decay as in Albert et al.^1^). The δ parameter is initially set to 0, and changes depending on error consistency according to:

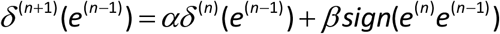

This equation indicates the following process: when two consecutive errors *e*^(n-1)^ and *e*^(n)^ have the same sign (i.e., they are consistent), the δ value that corresponds to the initial error *e*^(n-1)^ is increased by some amount β. On the other hand, the δ value decreases when the consecutive errors have opposite sign (i.e., they are inconsistent). Lastly, δ tends to decay towards 0 at a rate determined by α.

Note that δ varies as a function of error; at the same time, some errors may have a high δ value, while for others it is small. Its value depends on the past error history: which errors were experienced and whether they were consistent or inconsistent. To enable δ to vary with error, we treated it as a piecewise function that is constant within 3° error windows. We divided the error space between -99° and 99° in 3° bins and gave each bin its own independent δ value. Each δ value was updated according to the equation above. Note that when a δ value was not engaged on a given trial (because its error was not experienced) it only exhibited decay (no increase or decrease based on the β parameter).

In Fig. 2B we fit this model to the zero-variance and high-variance group simultaneously. The process we used was similar to that reported in our paper (see Ref. 4). Essentially, we used the error sequences that participants actually experienced to constrain the model. In other words, for a given model parameter set {α, β, *c, d*}, we simulated how error sensitivity would change with error and with trial number using errors experienced by the 13 zero-variance participants and the 12 high-variance participants. We then averaged the predicted error sensitivities across all participants. Because this process used noisy data from only 13 or 12 participants, it generated a noisy error sensitivity curve. For the purpose of model fitting, we needed to smooth this curve. We used a similar smoothing process as in Albert et al. For the zero-variance group we fit an exponential to the initial 20 cycles, and then a smoothing cubic splines over the remaining cycles (these functions were constrained to intersect on cycle 20). For the high-variance group, predicted error sensitivity curves were captured well by an exponential curve alone. We then input this smoothed error sensitivity curve into the state-space model in Section 1 to predict subject behavior. Note that the zero-variance and high-variance predicted curves were generated with the same model parameter set (in other words, subjects were treated as having the same baseline error sensitivity curve, and the same memory of errors update process: they only differed based on the errors they experienced). The model was fit by minimizing squared error between its predicted reach angles and zero-variance and high-variance reach angles simultaneously. Fig. 2B shows the best model’s predicted behavior. Supplementary Figure 2 shows model error sensitivities at different points during adaptation (cycles 1, 5, 40, 60). Note that combining the memory of errors (Albert et al.) with a natural error sensitivity versus error size decline (Wang et al.), closely captured the learning dynamics in Fig. 2B, and also predicted the timecourse by which behavior diverged across the variance conditions (Fig. 2C).

## Author contributions

STA analyzed data. STA and RS wrote manuscript.

## Data collection

146 volunteers (73 Male, 73 Female) aged 18-62 participated in our experiments. All experiments were approved by the Institutional Review Board at the Johns Hopkins School of Medicine. Informed consent was obtained. Participants were compensated $15/hour for participating in these studies.

## Data availability statement

The data were previously described in Ref. 4 and are deposited in OSF under accession code w5pkb.

## Code availability

Code for the state-space memory of errors model is deposited in OSF under accession code w5pkb.

